# Temperature fluctuations between years predict temporal allele frequency variation in a hybrid ant population

**DOI:** 10.1101/755496

**Authors:** R Martin-Roy, J Kulmuni

## Abstract

Genetic variability is essential for adaptation and isolated populations lacking variability can face extinction when environments change. A currently underappreciated source of genetic variability is hybridization with a closely related lineage. Hybridization can indeed facilitate adaptation e.g. via exchange of adaptive genetic material between species. Here we investigate the potential role of temperature in shaping the genetic variability in a hybrid population between *Formica polyctena* and *F. aquilonia* wood ants using long term population genetic data. We find that the frequencies of both parental-like alleles in the hybrid population co-vary with temperature over the years in males but not in females. Results suggest hybridization can lead to sex-specific outcomes that are dependent on ecological factors and that genetic variability resulting from hybridization could help hybrids cope with varying temperature.

## Introduction

Genetic variability is essential for adaptation to occur. It can rely on standing genetic variation (Barrett and Schluter 2008), de novo mutations (Orr 2005) or a third, to date understudied phenomenon: hybridization (i.e. interbreeding between two divergent lineages). Hybridization can increase genetic variation within a population much faster than new mutations and it has proven to be more common than previously thought. Over ten years ago it was estimated that 10% of animals and 25% of plants species hybridize (Mallet 2005), but the growing number of genomic studies highlighting cases of hybridization suggest that it could be even more common (Abbott et al. 2013; Mallet et al. 2016; Schumer et al. 2014; Taylor and Larson 2019). Furthermore, rates of hybridization are estimated to further increase due to climate change as species expand their ranges and new contact zones between closely related taxa will be formed (Chunco 2014).

Previously hybridization was thought to have mainly negative consequences for populations because of Dobzhansky-Muller incompatibilities (DMIs) (Dobzhansky 1936; Muller 1942). Such DMIs arise when alleles fixed in two distinct lineages occur within the same genome after interbreeding, creating negative epistatic interactions between alleles and leading to hybrid mortality or sterility (Orr 1996). However, it is now accepted that hybridization can also fuel adaptation via the creation of advantageous genetic combinations (Seehausen 2004; Grant et al. 2005; Abbott et al. 2013; Meier et al. 2017). Hybridization can therefore have an important role in evolution, by increasing genetic variation and providing novel phenotypes which could help adapt to new environments (Seehausen 2004; Grant et al. 2005; Abbott et al. 2013). Adaptive introgression has occurred for instance in *Heliconius* neotropical butterflies, where closely related species have exchanged protective colour patterning genes (The *Heliconius* Genome Consortium 2012). In some cases, hybridization can also fuel adaptive radiations and lead to the formation of new species e.g. in Darwin’s finches (Lamichhaney et al. 2018). As hybridization is common and can have major impacts on short- and long-term evolution, we need to understand its consequences in natural populations. However, only few long-term studies address the consequences of hybridization through time (but see Darwin’s finches; Grant, 2002 and desert sunflowers; Rieseberg et al., 2007).

Hymenoptera are excellent models for the study of hybridization because of their haplodiploidy. Males have a haploid genome and develop from unfertilized eggs while females (queens and workers) are diploid and arise through sexual reproduction. Consequently, recessive incompatibilities can be masked when heterozygous in females because of diploidy while they will be revealed in haploid males, where they can lead to hybrid breakdown (Koevoets and Beukeboom 2009; Kulmuni et al. 2010; Kulmuni and Pamilo 2014; Beukeboom et al. 2015). Similarly, recessive beneficial alleles can be selected for in haploid males more efficiently. The opportunity for long-term studies is also another reason why Hymenoptera are excellent models for studying hybridization. Location of wood ant nests can be stable over several tens of years, making it easy to monitor and collect genetic material over years. Previous studies have used hybridizing wood ants of the *Formica rufa* group to understand the consequences of hybridization in haplodiploids (Kulmuni et al. 2010; Kulmuni and Pamilo 2014). This group has multiple closely related species that have been estimated to diverge, partly in allopatry, during the last 490k years (Stockan and Robinson 2016). Currently, *Formica polyctena* has a more southern range while *F. aquilonia* has a boreo-alpine range (Fig. 1) (Stockan and Robinson 2016), but their distributions overlap in Southern Finland where multiple hybrid populations occur (Beresford et al. 2017). In this region, the Långholmen population encompasses more than 25 ant nests and is the most extensively studied hybrid population. Previous studies indicating the coexistence of two distinct genetic lineages named *F. polyctena*-like (formerly named R) and *F. aquilonia*-like (formerly named W) (Beresford et al. 2017; Kulmuni et al. 2019; Kulmuni and Pamilo 2014; Kulmuni et al. 2010). Both lineages are hybrids but currently inhabit different nests within the population (Kulmuni et al. 2010). Kulmuni & Pamilo (2014) have shown using microsatellite markers that sex-antagonistic selection acts on *F. polyctena*-like and *F. aquilonia*-like lineages: haploid males (especially from the *F. polyctena*-like lineage) suffer from hybrid breakdown during development when carrying alleles from the other lineage (i.e. introgressed alleles), while diploid females with introgressed alleles show hybrid vigour. Interestingly, while in the initial study alleles introgressed from *F. polyctena*-like lineage were not found in adult *F. aquilonia*-like males in 2004, they were detected at intermediate to high frequencies in 2014 (Kulmuni et al. 2019). Moreover, while in the initial microsatellite study *F. polyctena*-like alleles introgressed to *F. aquilonia*-like males were weakly selected against during development, a recent study performed with SNP markers (Kulmuni et al. 2019) showed selection favouring introgressed alleles over development in *F. aquilonia*-like males. These observations raise the question of what is causing the putative signal of fluctuating allele frequencies in the hybrid population over years. One potentially important variable fluctuating across years is temperature during development, which could cause differential survival of individuals depending on their genotypes.

**Figure 1:**
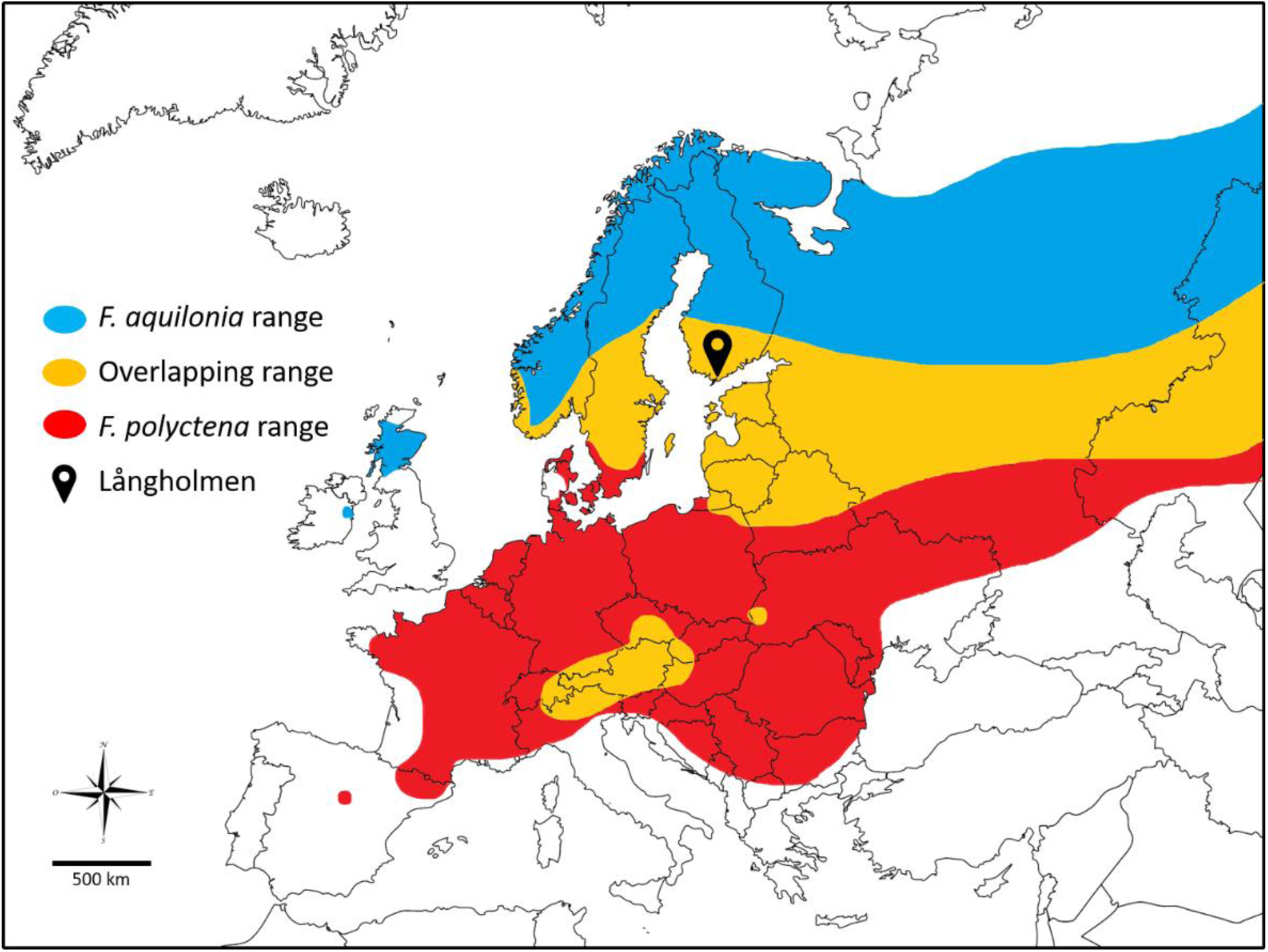
Map of *F. aquilonia* (blue) and *F. polyctena* (red) ranges. Overlapping ranges between the two species is represented in orange (adapted from Stockan and Robinson 2016).

In this study we investigate the role of temperature in shaping genetic variability of a hybrid ant population using long-term genetic data spanning 14 years. First, we ask if the frequency of *F. aquilonia*-like and *F. polyctena*-like alleles in the Långholmen hybrid population correlates with spring temperature over multiple years. Since the two hybridizing species have different distributions, northernmost for *F. aquilonia* and southernmost for *F. polyctena* (Fig. 1), we hypothesize they could be adapted to different temperatures, *F. aquilonia* being adapted to colder and *F. polyctena* to warmer temperatures. As a result, in the hybrid population, *F. polyctena*-like alleles would be selected for in warm years, leading to higher frequencies of *F. polyctena*-like alleles in warm years. Likewise, the frequency of *F. aquilonia*-like alleles should be higher in cold years. Second, we ask if the allele frequency variation across years displays a sex-specific pattern. This could be expected due to difference in ploidy between males and females. If selection acts on recessive alleles in haploid males but these are masked in heterozygous females, this would cause more frequency variation in haploid males compared to diploid females. Third, we ask if data from a single year supports the hypothesis that temperature during development causes differential survival of hybrid males with different parental alleles. Correlation between yearly spring temperature and parental allele frequency changes in hybrids could suggest context-dependent consequences of hybridization. This would also add evidence that genetic variation arising through hybridization could help populations adapt into changing temperatures (e.g. Jones et al. 2018; Oziolor et al. 2019).

## Material and Methods

### Samples

Here we study the *F. aquilonia* × *F. polyctena* hybrid population in Långholmen, Southern Finland. Previous work using microsatellite markers has identified two genetic lineages (*F. polyctena*-like and *F. aquilonia*-like) within the population, with lineage-specific (i.e. diagnostic) alleles and alleles introgressed between the lineages (Kulmuni et al. 2010; Kulmuni and Pamilo 2014). To assess if allele frequencies at these microsatellite loci co-vary with the average temperature during developmental season over multiple years, we collected and analysed samples from multiple time points. Parental species of the hybrid population are polygynous, having up to hundreds of queens within a mound (nest) and supercolonial, forming networks of interconnected mounds. The high number of reproductive queens in each nest induces average relatedness of near zero between individuals within a same nest. Consistent with this, no substructure between nests was previously observed within a lineage in the Långholmen hybrid population (see Supplementary from Kulmuni et al. 2019). Individuals from the same nest or from different nests within a population can therefore be considered as equally independent samples. In these species, queens mate once in their life, can live at least 5 years and have overlapping generations (Keller and Genoud 1997), therefore allelic combinations from a single queen can be subjected to selection over multiple years. Genotype data for adult individuals for the years 1996, 2004, 2008 and 2011 were obtained from Kulmuni et al. (2010) and Kulmuni and Pamilo (2014). In addition, ants collected in 2014 and 2018 were genotyped for the present study (see Supplementary material Table 1. for all samples used in this study).

To test if temperature during development could cause differential survival of hybrid males with different parental alleles, we utilized data from a single warm year (see results) and genotyped 42 males at early (small larva) stage and 29 males at late (adult) stage.

### Genotyping

DNA was extracted using the DNeasy Tissue kit (Qiagen) following manufacturer instructions. Genotyping was done using nine previously optimized microsatellite markers (Kulmuni & Pamilo 2014) shown to segregate independent of each other (no linkage disequilibrium). These markers were: FE7, FE13, FE19, FE17 (Gyllenstrand et al. 2002), FL29 (Chapuisat 1996) and, FY12, FY13, FY15 and FY3 (Hasegawa and Imai 2004). The loci were amplified by polymerase chain reaction (PCR) (Veriti 96 well Thermal Cycler; Applied Biosystems) with fluorescent labelling allowing for multiplexing markers with different dyes (i.e. three markers combined in one PCR). Type-it Microsatellites PCR kit (Qiagen) was used for master mix. Genotypes were resolved by capillary electrophoresis (3730 DNA Analyzer; Applied Biosystems) and scored with GeneMapper v.4.0 software (Applied Biosystems).

To control for potential bias due to samples being genotyped at different time points, 3 reference samples (i.e. samples of known genotype) were genotyped as control samples in each PCR. The final dataset after filtering low quality samples contained 787 individuals.

### Statistical analyses

To verify there is no nest effect on the genotype data, principal component analysis (PCA) (see Supplementary material Figure 1.) was ran to detect any nest substructure within a lineage using the *indpca* function from the *hierfstat* package (Goudet 2005) in R (R Core Team 2019, software v. 3.6). The following analyses were performed for each lineage and sex separately. Allele frequencies were calculated using the *pop.freq* function of the *hierfstat* package and transformed using the arcsine square root transformation. The allele status (i.e. introgressed *F. aquilonia*-like, introgressed *F. polyctena*-like, diagnostic *F. aquilonia*-like, diagnostic *F. polyctena*-like or origin unknown) was determined following Kulmuni et al. (2010). Temperature data from 1996 to 2018 were downloaded from the Finnish meteorological institute (https://en.ilmatieteenlaitos.fi, last accessed 13/08/2019). For each year, we used the mean temperature in April, as this is the time when majority of the new queens and males develop. Temperature data was collected from the Hanko Tvärminne meteorological station, located within a 500-meter radius of the study population. Then we asked if the change in allele frequencies between years correlates with temperature change between years. Allele frequency change was calculated by subtracting frequencies of a given year by the frequencies of the previous year (i.e. frequency delta = frequency year (n+1) – frequency year (n)) and similarly for temperature change. Change in allele frequencies was used as a response variable in a Linear Mixed Model, using the *lmer* function in the *lme4* package (Bates et al. 2015) assuming Gaussian distribution. *Temperature delta* and *locus* were used as fixed factors in the model and *year* was used as a random effect. *P*-values were calculated using *t* values assuming a normal distribution, the significance threshold being fixed at 5%. We report *R*^2^ for information only. Finally, we utilized data from a warm year to test if the distribution of parental alleles per individual was different between early and late developmental stages in males, as we would expect if temperature causes differential survival and fluctuations in male allele frequencies across years. To test for changes in the distribution of *F. aquilonia*-like diagnostic and *F. polyctena*-like introgressed alleles during development, we counted the number of these alleles present at the nine genotyped loci per single male in the *F. aquilonia*-like lineage, for both early (N = 42) and late (N = 29) developmental stages. Then, we performed a one-sided Wilcoxon rank sum test with continuity correction to estimate if the proportion of these alleles changed during male development according to our predictions (i.e. positive selection acting on *F. polyctena*-like alleles and negative selection acting on *F. aquilonia*-like alleles, since it was a warm year).

## Results

*F. aquilonia* has a boreo-alpine range (Stockan and Robinson 2016) (Fig. 1) and so alleles coming from this species are expected to be adapted to colder temperatures. Therefore, frequencies of *F. aquilonia*-like alleles were hypothesized to be higher during colder years potentially due to selection favouring individuals carrying these alleles at colder temperature. Alternatively, *F. polyctena* has a more Southern range (Stockan and Robinson 2016) (Fig. 1) and is expected to be adapted to warmer temperatures, compared to *F. aquilonia*. Consequently, their allele frequencies were expected to be higher in warmer years.

### *F. aquilonia*-like alleles have higher frequencies in males at colder temperatures

Year-to-year changes in allele frequencies (Frequency delta) and mean temperatures in April (Temperature delta) were plotted for each sex for *F. aquilonia*-like alleles, which correspond to both (1) *F. aquilonia*-like diagnostic alleles in the *F. aquilonia*-like lineage and (2) *F. aquilonia*-like introgressed alleles in the *F. polyctena*-like lineage (Fig. 2). Since there is no introgression in *F. polyctena*-like lineage males, these alleles were not plotted. As hypothesized, changes in allele frequencies behave differently between sexes (Fig. 2). An increase in mean temperature in April from one year to the next is coupled with a significant decrease in *F. aquilonia*-like diagnostic allele frequencies in males of the *F. aquilonia*-like lineage compared to the previous year (Estimate = −0.1229; SD = 0.2995; *p*-value < 0.0001, Fig. 2). For females, no significant correlation between change in allele frequencies and change in temperature of both *F. aquilonia*-like diagnostic and introgressed alleles was found.

**Figure 2:**
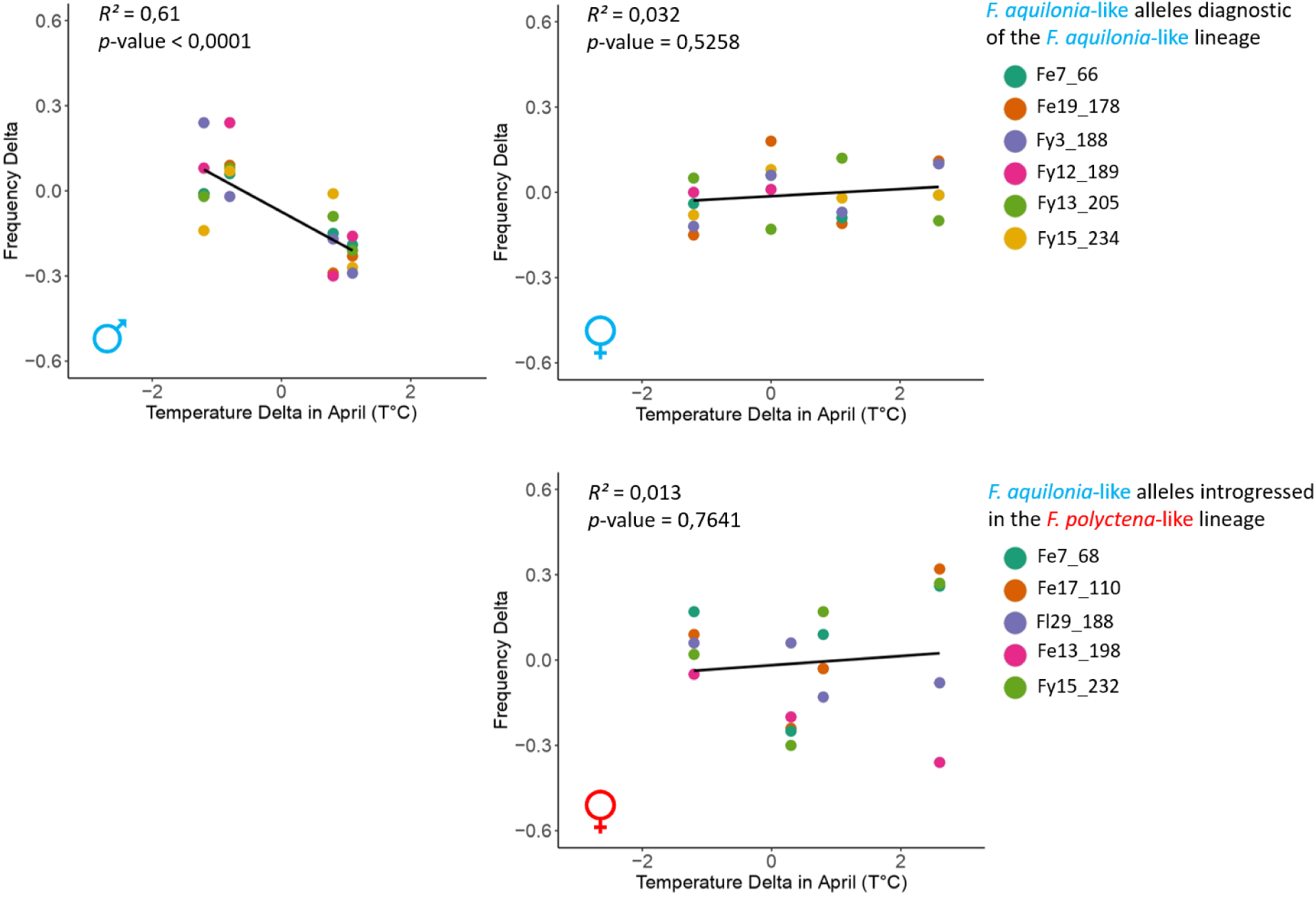
Year-to-year change in *F. aquilonia*-like allele frequency and mean temperature in April for *F. aquilonia*-like males and females of both lineages for introgressed and diagnostic alleles. *F. polyctena*-like males are not shown here because they never possess alleles introgressed from *F. aquilonia*-like lineage.

### *F. polyctena*-like alleles have higher frequencies in males at warmer temperatures

Similarly, year-to-year changes in allele frequencies and April temperature were plotted per sex for *F. polyctena*-like alleles, which correspond to both (1) *F. polyctena*-like diagnostic alleles of the *F. polyctena*-like lineage and (2) *F. polyctena*-like alleles introgressed in the *F. aquilonia*-like lineage (Fig. 3). *F. polyctena*-like diagnostic allele frequencies in both males and females were stable across years independently of mean temperature shifts (Fig. 3). However, in *F. aquilonia*-like males, when the temperature increased between years, introgressed alleles from the *F. polyctena*-like lineage significantly increased in frequency (Estimate = 0.3128; SD = 0.0682; *p*-value < 0.0001) (Fig. 3, lower left panel), as expected.

**Figure 3:**
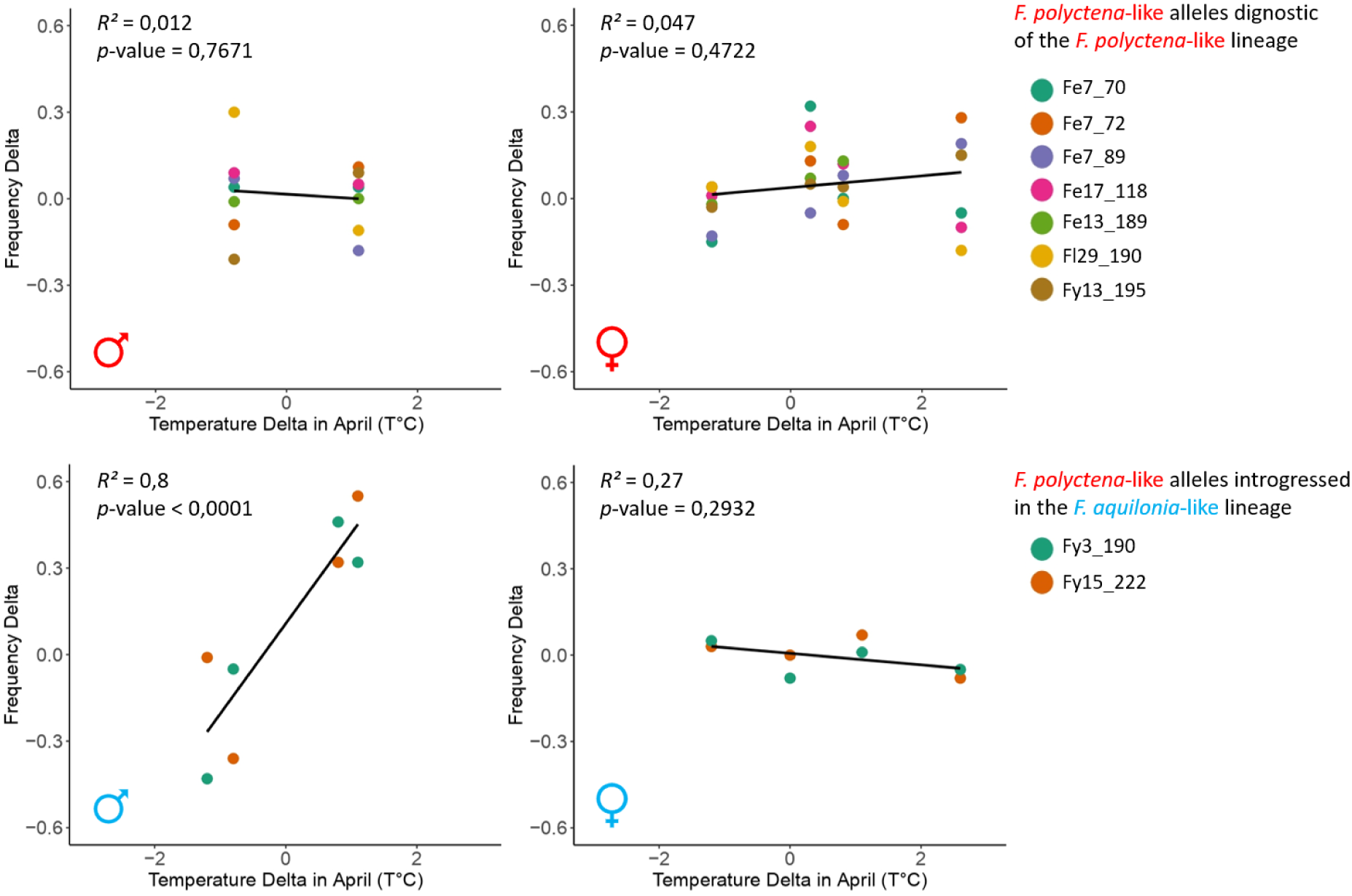
Year-to-year change in *F. polyctena*-like allele frequency and mean temperature in April for males and females of both lineages for introgressed and diagnostic alleles.

These results follow our expectations, namely that *F. aquilonia*-like alleles have higher frequencies at colder temperature and *F. polyctena*-like alleles have higher frequencies at warmer temperatures. To test if temperature-dependent selection during development could be responsible for frequency changes between years we measured larva to adult survival within a single year. We genotyped larvae and adults for the year 2014, which was a warm year (+2 °C compared to the 1963-2019 average, see Supplementary material Figure 2.). Therefore, we expected selection against diagnostic *F. aquilonia*-like alleles (putatively cold adapted) but selection for *F. polyctena*-like introgressed alleles (putatively warm adapted) in males during development. Comparing allele distributions at early and late developmental stage revealed a significant decrease in the number of diagnostic *F. aquilonia*-like alleles present in males (W = 839; *p*-value < 0.001) and a significant increase of *F. polyctena*-like introgressed alleles (W = 460.5; *p*-value = 0.025; Fig. 4), following our expectations for a warm year. Note that two out of nine loci have both diagnostic and introgressed alleles, and in this case the alleles are not independently selected.

**Figure 4:**
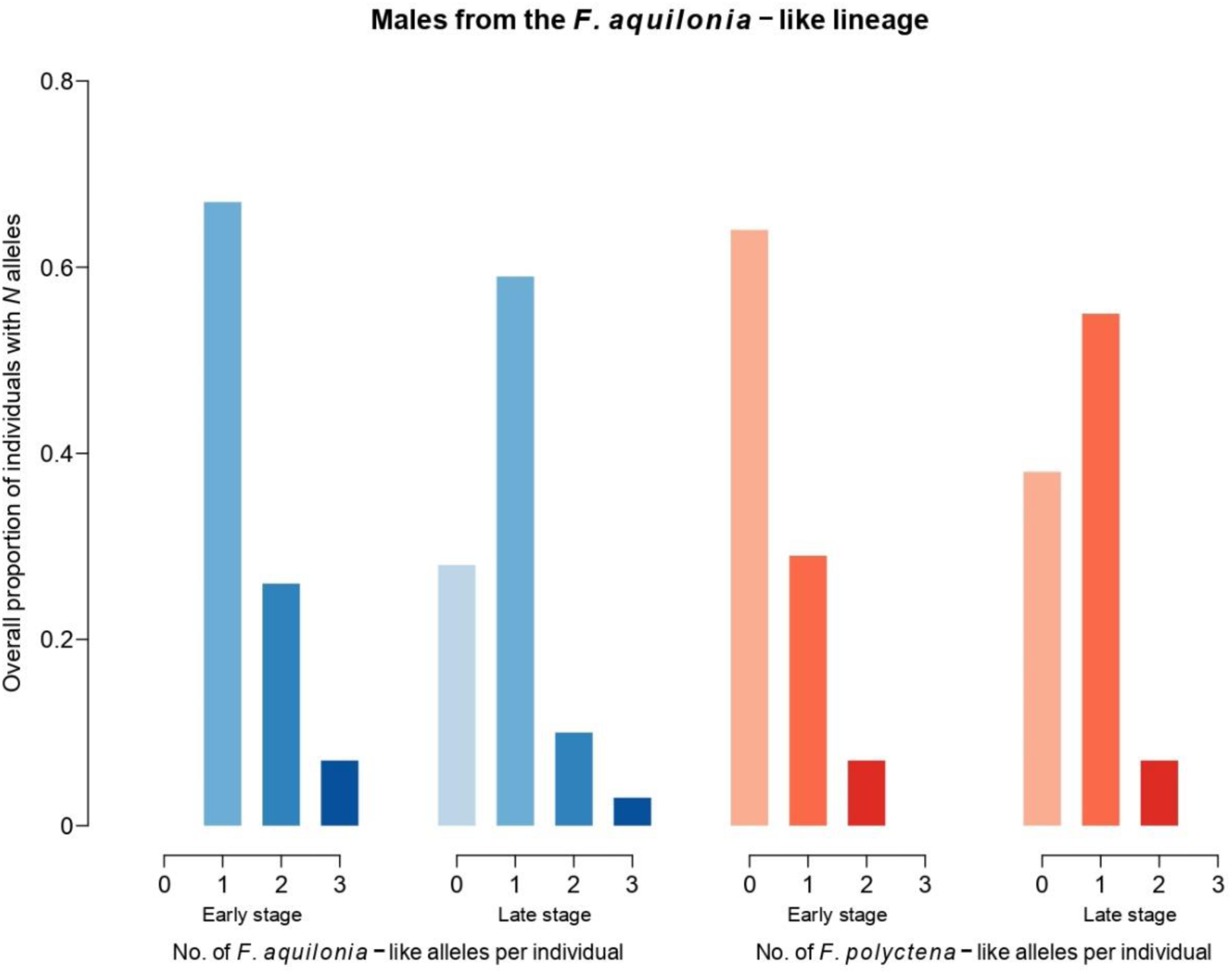
Proportion of males from the *F. aquilonia*-like lineage carrying *N* (from 0 up to 3) *F. aquilonia*-like diagnostic and/or *F. polyctena*-like introgressed alleles at the 9 genotyped microsatellite loci, for both early (N = 42) and late (N = 29) developmental stages, for the year 2014.

## Discussion

In order to survive, species and populations need to adapt to rapidly changing environments. Hybridization has been shown to increase genetic variability, which is essential for adaptation to occur. Accordingly, recent findings suggest hybridization could fuel rapid adaptation (e.g. Oziolor et al. 2019). This study aimed to understand if genetic variation within a hybrid population of *Formica* ants could be explained by temperature variation across years using long-term genetic data. To do so, we asked if allele frequency changes at nine loci were correlated with changes in the mean temperature in April (i.e. during sexual larvae development) over several years. Results show that in males, alleles from the Northernmost parental species have higher frequencies on cold years and alleles from the Southernmost parental species have higher frequencies on warm years. Furthermore, and in agreement with these results, we found that during a warm year, the number of alleles from the Southernmost parental species increased over development in males, while those of the Northernmost parental species decreased. This suggests temperature variation across multiple years could maintain (potentially adaptive) genetic diversity from both parental species within the hybrid population. These results show that outcomes of hybridization can be context-dependent and vary in time.

### Sex-specific allele frequency variation and temperature-dependent selection in a hybrid population

We investigated genetic variation over a 14-year period in a hybrid population between parental species with Northern (*F. aquilonia*) and Southern (*F. polyctena*) distributions. The results revealed significant correlation between allele frequency and temperature changes that was sex-specific. Indeed, in males, when temperature during development is warmer, *F. polyctena*-like alleles have higher frequencies. On the other hand, when temperature is colder, *F. aquilonia*-like alleles have higher frequencies. Furthermore, samples collected from the natural population in year 2014 revealed that *F. aquilonia*-like diagnostic alleles decreased significantly from early to late developmental stages while *F. polyctena*-like introgressed alleles increase, as expected on a warm year such as 2014. This result suggests temperature-dependent selection could drive allele frequency variation in males across years. However, under this scenario, linked selection on *F. aquilonia*-like diagnostic alleles could induce a frequency shift of *F. polyctena*-like introgressed alleles when they occur at the same microsatellite loci. Furthermore, it is unlikely that microsatellite alleles themselves are under selection, but they are near the true genomic targets of selection. Allele frequency change during development needs to be monitored over multiple years to confirm that thermal selection drives allele frequency variation across years in this hybrid ant population. Regarding females, allele frequency change between years did not show any correlation with temperature variation between years during development. In other taxa, the abiotic environment, particularly temperature, was shown to impact genetic diversity via frequency change of alleles under selection (e.g. Bergland et al. 2014; Bradbury et al. 2010; Dahlhoff and Rank 2007; Hornoy et al. 2015; Jump et al. 2006). However, genetic changes in the reproducing individuals of the population (mainly fathers) may not be as stark as suggested by the yearly variation, because ant queens can live multiple years and nests have overlapping generations.

Haldane’s rule predicts that when two species hybridize, it is the heterogametic sex that will suffer from hybrid breakdown (Haldane 1922). In haplodiploids, males are therefore expected to suffer from hybrid breakdown if they possess a deleterious allele. However, our results show that depending on abiotic factors, hybrid males will also experience positive selection upon having the correct allelic combination. The fact that female allele frequencies do not correlate with temperature changes could suggest that selection is more efficient in haploid males, while selection pressure is masked in diploid females (Beukeboom et al. 2015).

Diploidy also allows females to carry both *F. polyctena*-like and *F. aquilonia*-like alleles simultaneously in heterozygotes. Heterozygous status in females should therefore be positively selected as it guarantees higher fitness regardless of the temperature if alleles are co-dominant. This hypothesis is supported by results from Kulmuni and Pamilo (2014), who showed that heterozygous females for introgressed alleles were indeed selected for in the Långholmen hybrid population.

Our results suggest that fitness consequences of introgressed alleles are not only intrinsic but could be dependent on ecological factors. Hybridization increases genetic variability and can have various outcomes on the evolution of species, from the fusion of species to the creation of new ones (Seehausen 2004). In some instances, the introgression of a part of the genome of one parental population to the other could reveal DMIs, leading to hybrid breakdown (e.g. Brideau et al., 2006; Stelkens et al., 2015). However, introgression could also be beneficial, i.e. adaptive introgression (Abbott et al. 2013). Adaptive introgression is expected to rely on ecological opportunities (Seehausen 2004; Meier et al. 2017) and our results suggest that introgressed alleles could have either positive or negative impacts depending to the temperature. While more work is needed in the hybrid ant system to show a link between genotype, fitness and temperature, this could highlight an unappreciated aspect of hybrid populations i.e. selection could fluctuate between years, and a genotype that was deleterious in one environment could be beneficial in another environment.

### Future of hybrids in warmer temperatures

Under our thermal adaptation hypothesis and warmer temperatures induced by climate change, we could expect *F. polyctena*-like alleles to confer adaptive advantage to hybrid males possessing them. Having such alleles could be a way of coping with a warmer environment. Hybridization can occur between differentially temperature-adapted species (Pereira et al. 2016), which can lead to changes in temperature tolerance (Pereira et al. 2014) and the colonization of novel habitats (Gompert et al. 2006). Albeit sparse, genetic data suggests pure *F. polyctena* populations are in minority in Finland (Beresford et al. 2017), and hybridization could have allowed them to colonize this far North in the first place (e.g. Krehenwinkel and Tautz 2013). Indeed, new genetic combinations created by hybridization allows for novelties and colonizing new niches (e.g. Rieseberg et al., 2007). However, if the temperature keeps rising, we hypothesize that it could threaten *F. aquilonia* in which case *F. polyctena* alleles could allow *F. aquilonia* populations to cope with new thermal conditions (i.e. evolutionary rescue, Gonzalez et al., 2012), maintaining and potentially expanding the area of hybridization. Eventually we can even expect the hybrids to shift North as temperature increases and *F. aquilonia* alleles become non-adaptive under warmer temperatures.

### Conclusions

Long-term studies on the outcome of hybridization events are rare (e.g. Rieseberg et al., 2007; Grant, 2002). However, *Formica* ants offer here a unique insight into evolutionary mechanisms taking place in hybrid populations under a climate change context. Furthermore, key characteristics of this model like haplodiploidy and polygyny (i.e. multiple reproducing queens within a nest) offer great advantages when studying hybridization. Indeed, haplodiploidy allow the detection of selection on recessive alleles that would be otherwise very hard to detect in diploid models (if not located on sexual chromosomes). As for polygyny, increasing the rate of hybridization events allows for more introgression opportunities and buffers the consequences of deleterious introgression, because the high number of reproductive queens will insure colony survival despite some queens producing unfit offspring. In conclusion, this study showed that hybridization between two species of wood ants has led to the creation of new, potentially adaptive, allelic combinations that could help to cope with fluctuating ecological conditions.

## References

Abbott, R., D. Albach, S. Ansell, J. W. Arntzen, S. J. E. Baird, N. Bierne, J. Boughman, A. Brelsford, A. Buerkle, R. Buggs, R. K. Butlin, U. Dieckmann, F. Eroukhmanoff, A. Grill, S. H. Cahan, et al. 2013. Hybridization and speciation. Journal of Evolutionary Biology 26:229–246.

Barrett, R., and D. Schluter. 2008. Adaptation from standing genetic variation. Trends in Ecology & Evolution 23:38–44.

Bates, D., M. Mächler, B. Bolker, and S. Walker. 2015. Fitting Linear Mixed-Effects Models Using lme4. J. Stat. Soft. 67.

Beresford, J., M. Elias, L. Pluckrose, L. Sundström, R. K. Butlin, P. Pamilo, and J. Kulmuni. 2017. Widespread hybridization within mound-building wood ants in Southern Finland results in cytonuclear mismatches and potential for sex-specific hybrid breakdown. Mol. Ecol. 26:4013–4026.

Bergland, A. O., E. L. Behrman, K. R. O’Brien, P. S. Schmidt, and D. A. Petrov. 2014. Genomic evidence of rapid and stable adaptive oscillations over seasonal time scales in *Drosophila*. PLoS Genet. 10:e1004775.

Beukeboom, L. W., T. Koevoets, H. E. Morales, S. Ferber, and L. van de Zande. 2015. Hybrid incompatibilities are affected by dominance and dosage in the haplodiploid wasp *Nasonia*. Front. Genet. 6:140.

Bradbury, I. R., S. Hubert, B. Higgins, T. Borza, S. Bowman, I. G. Paterson, P. V. R. Snelgrove, C. J. Morris, R. S. Gregory, D. C. Hardie, J. A. Hutchings, D. E. Ruzzante, C. T. Taggart, and P. Bentzen. 2010. Parallel adaptive evolution of Atlantic cod on both sides of the Atlantic Ocean in response to temperature. Proc. R. Soc. B 277:3725–3734.

Brideau, N. J., H. A. Flores, J. Wang, S. Maheshwari, X. Wang, and D. A. Barbash. 2006. Two Dobzhansky-Muller genes interact to cause hybrid lethality in *Drosophila*. Science 314:1292–1295.

Chapuisat, M. 1996. Characterization of microsatellite loci in *Formica lugubris* B and their variability in other ant species. Mol. Ecol. 5:599–601.

Chunco, A. J. 2014. Hybridization in a warmer world. Ecol. Evol. 4:2019–2031.

Dahlhoff, E. P., and N. E. Rank. 2007. The role of stress proteins in responses of a montane willow leaf beetle to environmental temperature variation. J. Biosci. 32:477–488.

Dobzhansky, T. 1936. Studies on hybrid sterility. II. Localization of sterility factors in *Drosophila pseudoobscura* hybrids. Genetics 21:113.

Gompert, Z., J. A. Fordyce, M. L. Forister, A. M. Shapiro, and C. C. Nice. 2006. Homoploid hybrid speciation in an extreme habitat. Science 314:1923–1925.

Gonzalez, A., O. Ronce, R. Ferriere, and M. E. Hochberg. 2012. Evolutionary rescue: an emerging focus at the intersection between ecology and evolution. Phil. Trans. R. Soc. B: Biological Sciences 368:20120404–20120404.

Goudet, J. 2005. hierfstat, a package for r to compute and test hierarchical F-statistics. Mol. Ecol. Notes 5:184–186.

Grant, P. R. 2002. Unpredictable Evolution in a 30-Year Study of Darwin’s Finches. Science 296:707–711.

Grant, P. R., B. R. Grant, and K. Petren. 2005. Hybridization in the Recent Past. The American Naturalist 166:56–67.

Gyllenstrand, N., P. J. Gertsch, and P. Pamilo. 2002. Polymorphic microsatellite DNA markers in the ant *Formica exsecta*. Mol. Ecol. Notes 2:67–69.

Haldane, J. B. S. 1922. Sex ratio and unisexual sterility in hybrid animals. Journal of Genetics 12:101–109.

Hasegawa, E., and S. Imai. 2004. Characterization of microsatellite loci in red wood ants *Formica* (s. str.) spp. and the related genus *Polyergus*. Mol. Ecol. Notes 4:200–203.

Hornoy, B., N. Pavy, S. Gérardi, J. Beaulieu, and J. Bousquet. 2015. Genetic adaptation to climate in white spruce involves small to moderate allele frequency shifts in functionally diverse genes. Genome Biology and Evolution 7:3269–3285.

Jones, M. R., L. S. Mills, P. C. Alves, C. M. Callahan, J. M. Alves, D. J. R. Lafferty, F. M. Jiggins, J. Jensen, J. Melo-Ferreira, and J. M. Good. 2018. Adaptive introgression underlies polymorphic seasonal camouflage in snowshoe hares. Science 360:1355–1358.

Jump, A. S., J. M. Hunt, J. A. Martínez-Izquierdo, and J. Peñuelas. 2006. Natural selection and climate change: temperature-linked spatial and temporal trends in gene frequency in *Fagus sylvatica*. Mol. Ecol. 15:3469–3480.

Keller, L., and M. Genoud. 1997. Extraordinary lifespans in ants: a test of evolutionary theories of ageing. Nature 389:958–960.

Koevoets, T., and L. W. Beukeboom. 2009. Genetics of postzygotic isolation and Haldane’s rule in haplodiploids. Heredity 102:16–23.

Krehenwinkel, H., and D. Tautz. 2013. Northern range expansion of European populations of the wasp spider *Argiope bruennichi* is associated with global warming-correlated genetic admixture and population-specific temperature adaptations. Mol. Ecol. 22:2232–2248.

Kulmuni, J., P. Nouhaud, L. Pluckrose, I. Satokangas, K. Dhaygude, and R. K. Butlin. 2019. Evidence for natural selection and barrier leakage in candidate loci underlying speciation in wood ants. BioRxiv 500116.

Kulmuni, J., and P. Pamilo. 2014. Introgression in hybrid ants is favored in females but selected against in males. Proceedings of the National Academy of Sciences 111:12805–12810.

Kulmuni, J., B. Seifert, and P. Pamilo. 2010. Segregation distortion causes large-scale differences between male and female genomes in hybrid ants. Proceedings of the National Academy of Sciences 107:7371–7376.

Lamichhaney, S., F. Han, M. T. Webster, L. Andersson, B. R. Grant, and P. R. Grant. 2018. Rapid hybrid speciation in Darwin’s finches. Science 359:224–228.

Mallet, J. 2005. Hybridization as an invasion of the genome. Trends in Ecology & Evolution 20:229–237.

Mallet, J., N. Besansky, and M. W. Hahn. 2016. How reticulated are species? BioEssays 38:140–149.

Meier, J. I., D. A. Marques, S. Mwaiko, C. E. Wagner, L. Excoffier, and O. Seehausen. 2017. Ancient hybridization fuels rapid cichlid fish adaptive radiations. Nat. Commun. 8:14363.

Muller, H. 1942. Isolating mechanisms, evolution, and temperature. Biol. Symp. 6:71–125.

Orr, H. A. 1996. Dobzhansky, Bateson, and the genetics of speciation. Genetics 144:1331.

Orr, H. A. 2005. The genetic theory of adaptation: a brief history. Nat. Rev. Genet. 6:119–127.

Oziolor, E. M., N. M. Reid, S. Yair, K. M. Lee, S. Guberman VerPloeg, P. C. Bruns, J. R. Shaw, A. Whitehead, and C. W. Matson. 2019. Adaptive introgression enables evolutionary rescue from extreme environmental pollution. Science 364:455–457.

Pereira, R. J., F. S. Barreto, and R. S. Burton. 2014. Ecological novelty by hybridization: experimental evidence for increased thermal tolerance by transgressive segregation in *Tigriopus californicus*: ecological novelty by hybridization. Evolution 68:204–215.

Pereira, R. J., I. Martínez-Solano, and D. Buckley. 2016. Hybridization during altitudinal range shifts: nuclear introgression leads to extensive cyto-nuclear discordance in the fire salamander. Mol. Ecol. 25:1551–1565.

Rieseberg, L. H., S.-C. Kim, R. A. Randell, K. D. Whitney, B. L. Gross, C. Lexer, and K. Clay. 2007. Hybridization and the colonization of novel habitats by annual sunflowers. Genetica 129:149–165.

Schumer, M., G. G. Rosenthal, and P. Andolfatto. 2014. How common is homoploid hybrid speciation? Evolution 68:1553–1560.

Seehausen, O. 2004. Hybridization and adaptive radiation. Trends in Ecology & Evolution 19:198–207.

Stelkens, R. B., C. Schmid, and O. Seehausen. 2015. Hybrid breakdown in cichlid fish. PLoS ONE 10:e0127207.

Stockan, J. A., and E. J. Robinson. 2016. Wood ant ecology and conservation. Cambridge University Press.

Taylor, S. A., and E. L. Larson. 2019. Insights from genomes into the evolutionary importance and prevalence of hybridization in nature. Nat. Ecol. Evol. 3:170–177.

The Heliconius Genome Consortium, K. K. Dasmahapatra, J. R. Walters, A. D. Briscoe, J. W. Davey, A. Whibley, N. J. Nadeau, A. V. Zimin, D. S. T. Hughes, L. C. Ferguson, S. H. Martin, et al. 2012. Butterfly genome reveals promiscuous exchange of mimicry adaptations among species. Nature 487:94–98.

