## Supplementary material for "Temperature fluctuations between years predict temporal allele frequency variation in a hybrid ant population"

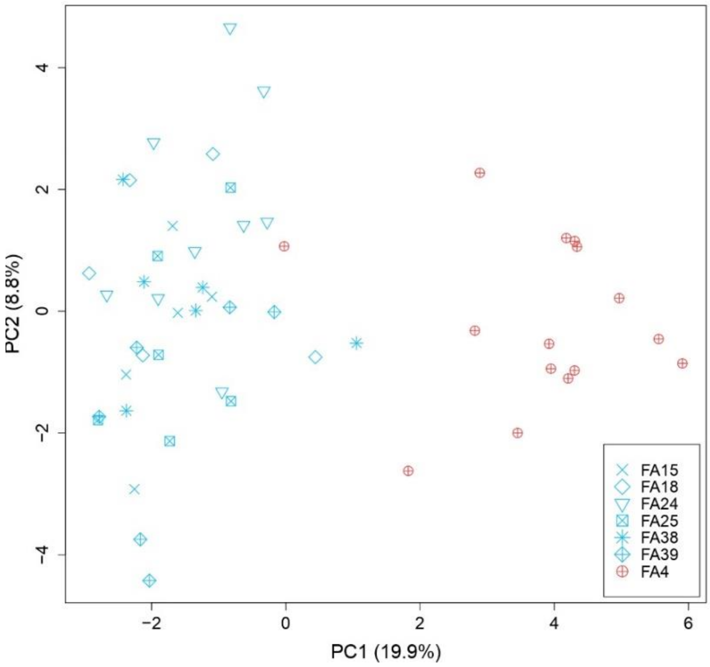


**Figure 1:** Principal component analysis of the Långholmen population, in blue is represented the *F. aquilonia*-like lineage and in red the *F. polyctena*-like lineage. Each nest is indicated by a symbol, each datapoint is an individual. The two lineages are separated in two distinct groups and no substructure due to a nest effect is present within the *F. aquilonia*-like group in blue.

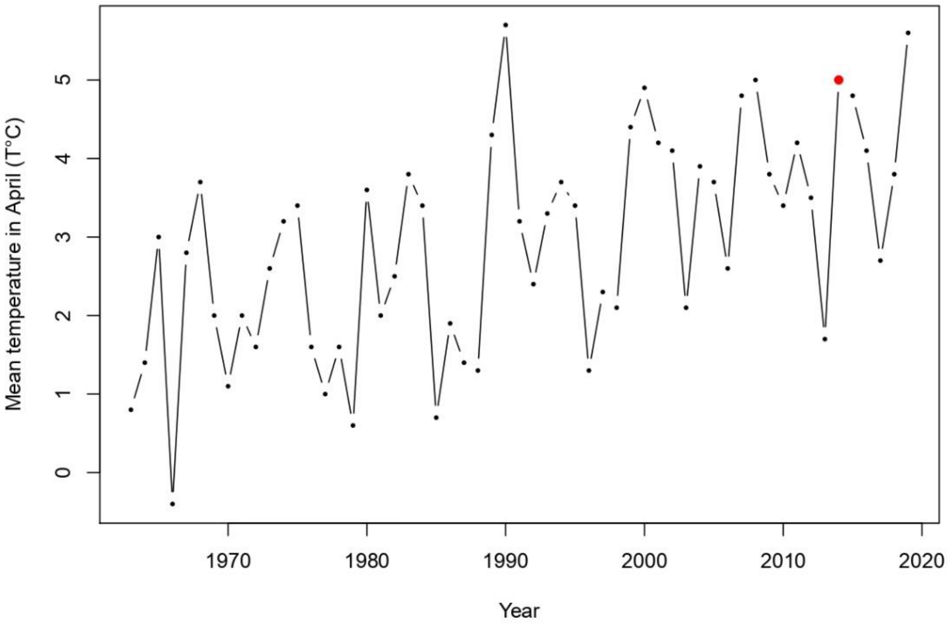


**Figure 2:** Mean April temperature at the Hanko Tvärminne station from 1963 to 2019, the 2014 year is represented in red.

| Population | Species identity | Mitochondrial DNA haplotype | Sex | 1996 | 2004 | 2008 | 2011 | 2014 | 2018 |
|---|---|---|---|---|---|---|---|---|---|
| Langholmen (*F. aquilonia* -like) | Hybrids | *F. polyctena* mtDNA | male | | N=62 | N=41 | N=28 | N=71 | N=38 |
| | | | female | N=80 | N=70 | N=100 | | N=31 | N=37 |
| Langholmen (*F. polyctena* -like) | Hybrids | *F. polyctena* mtDNA | male | | N=35 | N=41 | N=23 | | |
| | | | female | N=9 | N=25 | | N=34 | N=32 | N=30 |

**Table 1:** Samples collected and genotyped from the Långholmen population over the years. The genetic groups of the Långholmen population and mitochondrial haplotypes (from Kulmuni et al. 2010) are also indicated.
